# Identification of the novel inhibitors against *M. tuberculosis* ESX-1 secretion system EccA_1_ enzyme using virtual screening, docking and dynamics simulation techniques

**DOI:** 10.64898/2026.04.09.717399

**Authors:** Ramesh Kumar, Ajay K. Saxena

**Affiliations:** Rm-403/440, Structural Biology Lab, School of Life Sciences, Jawaharlal Nehru University, New Dehi-110067

**Keywords:** *M. tuberculosis* ESX-1 secretion system, EccA1, ADP, ZINC compounds, Drugs

## Abstract

The *M. tuberculosis* ESX-1 secretion system EccA_1_ enzyme is involved in the secretion of virulence factors and is essential for virulence and bacterial survival within the phagosome. Development of the small molecular inhibitors abolishing EccA_1_ function can yield new antivirulence drugs. In this study, we modeled the full-length EccA_1_ (573 residues, Mw ∼62.4 kDa) structure, which contains N-terminal TPR domain and a C-terminal CbxX/CfqX type ATPase domain. We have identified five ZINC compounds having binding energy i. e. Z1 (ZINC000004513760, -43.45 kcal/mol), Z2 (ZINC000000001793, -49.56 kcal/mol), Z3 (ZINC000005390388, -55.83 kcal/mol), Z4 (ZINC000257294577, -52.33 kcal/mol), Z5 (ZINC000004824264, -44.44 kcal/mol) through virtual screening of the ZINC compounds targeting C-terminal ATPase pocket of EccA_1_. The Z1-Z5 compounds were compared with ADP substrate having binding energy (Adenosine diphosphate, -35.00 kcal/mol), p97 ATPase inhibitors i.e. NMS873 (3-[3-cyclopentylsulfanyl-5-[[3-methyl-4-(4 methylsulfonylphenyl)phenoxy]methyl]-1,2,4-triazol-4-yl]pyridine, -48.68 kcal/mol), and CB5083 (1-[4-(benzylamino)-5H,7H,8H-pyrano[4,3-d]pyrimidin-2-yl]-2-methyl-1H-indole-4-carboxamide, -50.88 kcal/mol) against EccA1. The Z1-Z5 compounds exhibited good Absorption, Distribution, Metabolism, and/or Excretion properties (ADMTE). Pharmacokinetic properties and Lipinsky’s rule of five for Z1-Z5 compounds showed drug-like properties. 100 ns dynamics simulation analysis on EccA_1_ complexed with (i) Z1-Z5 compounds (ii) ADP substrate and (iii) NMS873 and CB5083 inhibitors showed high stability and biologically relevant conformation during dynamics simulation. These data indicate that Z1-Z5 compounds may act as potential inhibitors against EccA_1_ and provide avenues for new antivirulence drug development after *in vitro* and *in vivo* clinical trials.

## Introduction

The World Health Organization (WHO) estimated 10.8 million people were infected with tuberculosis worldwide in 2023, and it caused 1.25 million deaths (∼161,000 people with HIV) in 2024 [1]. The MDR and XDR-TB remain a global health issue, and ∼25,000 to 30,000 XDR-TB cases were detected in 2023 worldwide [1]. WHO has recommended isoniazid, rifampicin, pyrazinamide, ethambutol, and streptomycin as first lines of the drugs, while ethionamide and BPaLM regimen for XDR and MDR-TB [2]. Beyond vaccines, a new class of small molecules targeting the key mycobacterial proteins needs to be developed to cure XDR and MDR-TB. The availability of small molecular databases and improved automation of the target-based drug discovery process have contributed to the new drug candidate development [2].

The *M. tuberculosis* EccA1 enzyme (573 residues, 63kDa) from the ESX-1 secretion system is essential for the secretion of ESX-1 virulence proteins for pathogenicity, and its inhibition will abolish bacterial virulence and its survival in the host [3]. The EccA1 contains an N-terminal TPR domain (∼30 kDa) and a C-terminal CbxX/CfqX type ATPase domain, which functions as the oligomerization domain and exhibits ATPase activity. The EccA1 enzyme is involved in EspAC virulence factor secretion, and knockout of the EccA1 blocks the virulence proteins secretion [4]. The EccA1 is unique to the pathogen and absent in the human and offers an attractive target for drug development. The EccA1 active site offers an attractive target to develop small molecular inhibitors for ATP binding and hydrolysis. The EccA1 is evolutionarily conserved and present across *M. tuberculosis* species, thus offering selectivity.

ATP competitive CB5083 specifically inhibits the D2 domain of AAA-ATPase p97 and is orally bioavailable [5]. Treatment of CB5083 to tumor cells led to significant impairment of the ubiquitin−proteasome system (UPS) and protein degradation, endoplasmic reticulum-associated degradation (ERAD) functions, which led to proteotoxic stress and cell death [6]. The CB5083 is undergoing Phase-1 clinical trials as a drug candidate for the treatment of cancer. The NMS873 is the most potent VCP inhibitor, interferes with autophagy, induces cancer cell death, and is involved in the activation of the unfolded protein response [7]. The NMS873 covalently modifies the active site Cys523 and blocks the ATP binding.

Traditional anti-TB drugs exhibit high potency against chronic mycobacterial infection. The virulence inhibitors against the ESX-1 system may pose low drug resistance developed by mycobacteria. Since EccA1 is essential for the ESX-1 virulence factor secretion/pathogenicity, its inhibition will reduce virulence and bacterial survival in the macrophage. In the current study, we have done molecular modeling of the EccA1, structure-based virtual screening of the EccA1, and validation of the discovered ZINC compounds using (i) molecular docking, (ii) binding free energy delta G, and (iii) assessment of the stability of the EccA1-inhibitor complex using the dynamics simulation technique. We have studied the pharmacokinetic and toxicity properties of the discovered ZINC compounds and compared them with the natural ADP substrate and two clinical inhibitors, the CB5083 and NMS873, against p97 AAA+ ATPase. The five lead ZINC compounds against EccA1 will provide avenues for future antivirulence drug development against *M. tuberculosis*.

## Materials and methods

### Structural modeling of the EccA1 enzyme

The EccA1 gene (1-573 residues, Mw∼62.4 kDa) was retrieved from Uniprot (P9WPH9) database and built the structure using Alpha-fold server [8]. PROCHECK [9] server was used for stereochemical validation and Ramachandran plot analysis of the EccA1 model. The Maestro from the SchroÖdinger program [10] and the LIGPLOT program [11] were used for protein-ligand interaction analysis.

### Protein preparation and grid generation

Protein Preparation Wizard [12] module of SchrÖdinger [10] was used for initial processing of the EccA1 model, which added the missing hydrogen atoms and assigned the bond ordering. Prime module of the SchrÖdinger was used to fill all missing side chains. Receptor grid generation wizard of the Glide module [13] was used for receptor grid box generation using Pro336, Thr338, Lys340, and Arg429 residues of the ATPase pocket of the EccA1.50,000 steps of the steepest minimization were performed on EccA1 model using OPLS-3 (optimized potentials for liquid simulations) force field [14]. The grid box was generated using the centroid of the ATPase pocket and optimized potentials for ligand docking.

### Compounds library screening

ZINC database library (drug-like compounds) [15] was screened against the ATPase pocket of EccA1 using the Maestro module [10] and yielded the top 5 potential ZINC compounds (Z1-Z5). Energy minimization was performed on discovered Z1-Z5 compounds using the OPLS-3 force field until RMSD ∼ reached 2.0 Å. LIGPREP module of SchrÖdinger was used to build the Z1-Z5 compounds, and GLIDE module in XP (extra precision) mode was used to dock the Z1-Z5 compounds in the ATPase pocket of EccA1.ADP ligand was also docked into the ATPase pocket of EccA1 to screen substrate binding into pocket. Two drugs like ATPase inhibitors NMS873 (Pub: 71521142) [16] and CB5083 (Pub: 97428) [17] were downloaded in SDF format and prepared by optimizing their geometries, assigning proper protonation states at physiological pH, and performing energy minimization using the LigPrep module of SchrÖdinger. GLIDE was used to dock two drugs in the ATPase pocket of the EccA1.The ADP substrate and two drugs NMS873 and CB5083 were used for comparative analysis with the discovered Z1-Z5 compounds.

### Binding energy calculation using Prime MM-GBSA

The MM-GB/SA (Prime molecular mechanics-generalized born surface area) module of the SchrÖdinger program [18] was used to calculate the free energy of the EccA1-ligand complexes. OPLS3 force field [19] and GB/SA (Generalized Born/Surface area) continuum 2.0 solvent model [20] were used for energy calculation of minimized EccA1-ligand complexes. The binding free energy (ΔG_bin_) was calculated using dynamics simulation trajectories of all EccA1-ligand complexes. The binding free energy depends on (i) gas-phase free energy (DGMM), (ii) the solvation free energy (ΔG_sol_), and (iii) the change in the system entropy (-TΔS). Molecular mechanics Poisson-Boltzmann Surface Area (MM-GBSA) module, as shown ΔGbin = ΔGMM + ΔGsol -TΔS was used to compute the binding free energies [21, 22]. The PLIP server was used to rationalize the dominating interactions in the protein-ligand system [23].

### Analysis of the ADME (Absorption, Distribution, Metabolism, and Excretion) properties

The QikProp module of the SchrÖdinger [24] was used to calculate the ADMET properties of the discovered ZINC compounds. Lipinski rule of five (No. of hydrogen bond donors and acceptors, Log P), dipole moment, molecular weight, and oral absorption parameters were calculated for all ZINC compounds. The 6-31G basis set in the Jaguar module of SchrÖdinger [25] was combined with the B3LYP hybrid function.

Prime MMGBSA (Molecular Mechanics Generalized Born Surface Area) [26], post-docking application of SchrÖdinger, was used to analyze the binding free energy of all ZINC compounds and ranked based on MM-GBSA scores. The ADMET properties and Lipinski’s rule of five will help in distinguishing between drug-like and non-drug-like features in the discovered compounds.

### Molecular dynamics simulation

The apo EccA1 and its complexes with (i) Z1-Z5 compounds (ii) ADP substrate, and (iii) NMS873 and CB5083 drugs were subjected to the 100 ns dynamics simulation. The Desmond module of SchrÖdinger 2019-4 [28] was used for dynamics simulation with explicit solvent OPLS3 force field. The crystallographic TIP3P water molecules under orthorhombic periodic boundary conditions for a 10 Å buffer region were used. Counterions were used to neutralize the charge of the whole system. A constant temperature (300 K) and pressure (1 bar) were maintained for the whole system using the NPT ensemble of Nose-Hoover thermostat [29] and barostat, respectively. Limited memory Broyden-Fletcher-Goldfarb-Shanno (LBFGS) algorithm [30] (Threshold∼1kcal/mol/Å) was used for energy minimization. Long-range electrostatic interactions (cut-off radius∼9 Å) and short-range Coulomb and van der Waals interactions were used. Trajectories were analyzed with the Maestro program [10] after collecting data every 100 ps.

### Post-dynamics simulation analysis

The radius of gyration (Rg), RMSF, RMSD, hydrogen bonding, and SASA were calculated for apo and EccA1 complexes with Z1-Z5 compounds, ADP, and NMS-873 & CB-5083 drugs.

## Results

### Structural features of the EccA1 enzyme

The EccA1 forms a hexameric ATPase and belongs to the CbxX/Cbfq family of enzymes, and hydrolyzes ATP weakly compared to its ATPase domain. Alpha-fold server [32] was used to build the structure of full-length EccA1(1-573 residues). The PROCHECK [33] analysis on the EccA1 model showed good stereochemistry, and most of the residues lie in the allowed region of the Ramachandran plot (Fig. S1). The EccA1 model contains an N-terminal TPR domain (1-272 residues, yellow), which consists of six tetratricopeptide repeats (TPR1-TPR6) and a short β-hairpin motif between TPR2 and TPR3 motifs (green). The β-hairpin motif of the TPR domain (Fig. 1B, 76-82 residues, green) binds to the export arm of the EspC virulence factor (Fig. 1B, magenta) and is involved in virulence factor secretion.

**Fig. 1.**
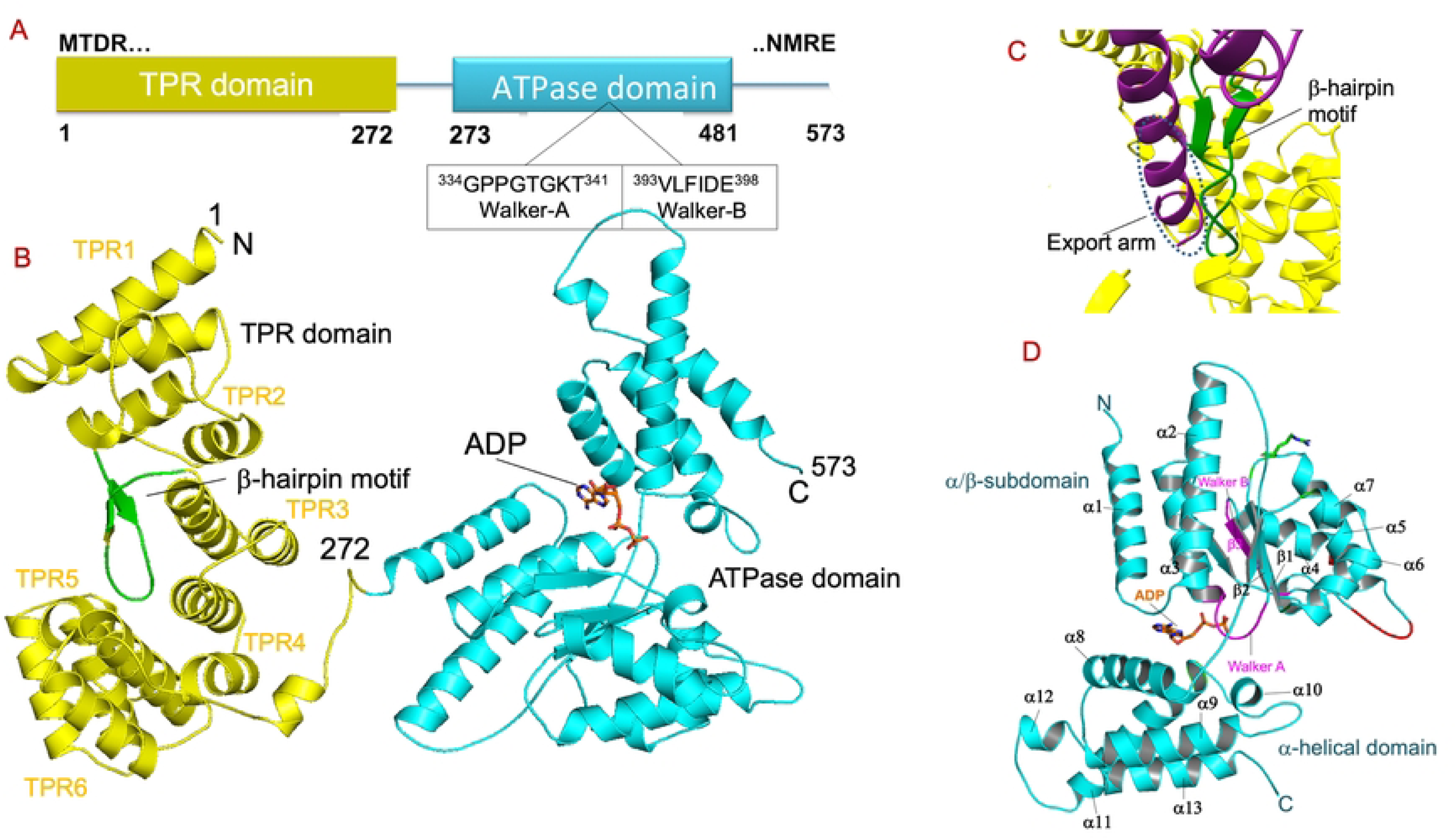
Overall architecture of *M. tuberculosis* EccA1. **(A**) Schematic diagram of the full length EccA1. The EccA1 consists of the N-terminal TPR domain (1-265 residues, yellow) and C-terminal ATPase domain (273-573 residues, cyan) connected by small linker 266-275. The Walker A (^334^GPPGTGKT^341^) and Walker-B (^393^VLFIDE^398^) motifs involved in ATP/Mg^2+^ binding is shown in the figure. **(B)** The stereoview of the full length EccA1. The N-terminal TPR domain (1-265 residues, yellow) consists of TPR1-TPR6 Trico-peptide repeat domains and a β-hairpin motif (green) between TPR2 and TPR3 motifs. The C-terminal ATPase domain is divided into large α/β helical subdomain (276-462, cyan) and a small α-helical subdomain (467-573, residues, cyan). The ADP binding sites is located at the interface between these two subdomains. **(C)** Figure showing the interaction between EspC export arm (magenta) and β-hairpin motif of the TPR domain (green). **(D)** Stereoview of the C-terminal ATPase domain of EccA1. The protein consists of α1-α13 helices, β1-β3 sheets (cyan), Walker A and Walker B motifs (pink), Arginine sensor (green) and ATP (orange).

The C-terminal ATPase domain (273-573 residues) of EccA1 contains an α/β subdomain and an α-helical domain (Fig. 1D, cyan), joined together with a short linker. The Walker-A and Walker-B motifs involved in ATP/Mg^2+^ binding is observed in the α/β subdomain of the EccA1. In the α/β subdomain, seven α-helices encircle the five β strands that are placed in a 2-3-6-1-7 arrangement at the core. Six α-helical bundles make up the minor α-helical domain. Nucleotide binding is carried out by the classical Walker A (^334^GPPGTGKT^341^) and Walker B (^393^VLFIDE^398^) motifs found in the ATPase pocket of the α/β subdomain. Our crystal structure analysis of the C-terminal ATPase domain complexed with ATP+Mg^2+^ showed the post-hydrolytic state of EccA1 bound with ADP only (data not shown) and used for comparative analysis with discovered ZINC compounds. The EccA1 is involved in mycolic acid synthesis and virulence factor secretion in *M. tuberculosis* [34]. The EccA1 is conserved in other mycobacterial species and is a potential target for antivirulence drugs development.

### Molecular docking and interactions analysis of the ADP substrate and Z1-Z5 inhibitors

The ATPase pocket of EccA1 is accessible for small molecular inhibitors, which need to penetrate the host macrophage, granuloma, and mycobacterial cell wall. To discover specific ATPase inhibitors, 7.5 million drug-like compounds from the ZINC database were virtually screened against the ATPase pocket of EccA1, which have drug-like properties. All ZINC compounds were docked to the predicted ATPase pocket, and potential compounds with the highest affinity towards EccA1 were selected for further analysis. The glide docking module of Schrödinger [10] employs advanced algorithms to predict the binding affinity and geometry of small compounds against target proteins. It aids in the identification of potential drug compounds efficiently and accurately.

Molecular docking result with ADP substrate showed the docking score -8.3 kcal/mol and binding affinity -35.00 kcal/mol. Five potential ZINC compounds were identified, having top ranks and showing the binding energy, e.g., Z1 (−43.45 kcal/mol), Z2 (−49.56 kcal/mol), Z3 (−55.83 kcal/mol), Z4 (−52.33 kcal/mol), and Z5 (−44.44 kcal/mol). The docked poses of ADP and five ZINC compounds (Z1-Z5) are shown at the ATPase pocket of EccA1 (Fig. 2A-F). The Z1-Z5 compounds showed better docking scores and the lowest MM-GBSA binding energy against EccA1, when compared to ADP (Table 1). The Z1-Z5 compounds showed high affinity due to their more lipophilic character and hydrogen bonding. The Z1-Z5 compounds contain a purine-like ring, a ribose sugar, and an extended acidic group, resembling to Adenosine or other ADP analogs. The 2D image of Z1-Z5 compounds showed quite similar structures to purine nucleosides, having nucleotide-binding properties.

**Fig. 2.**
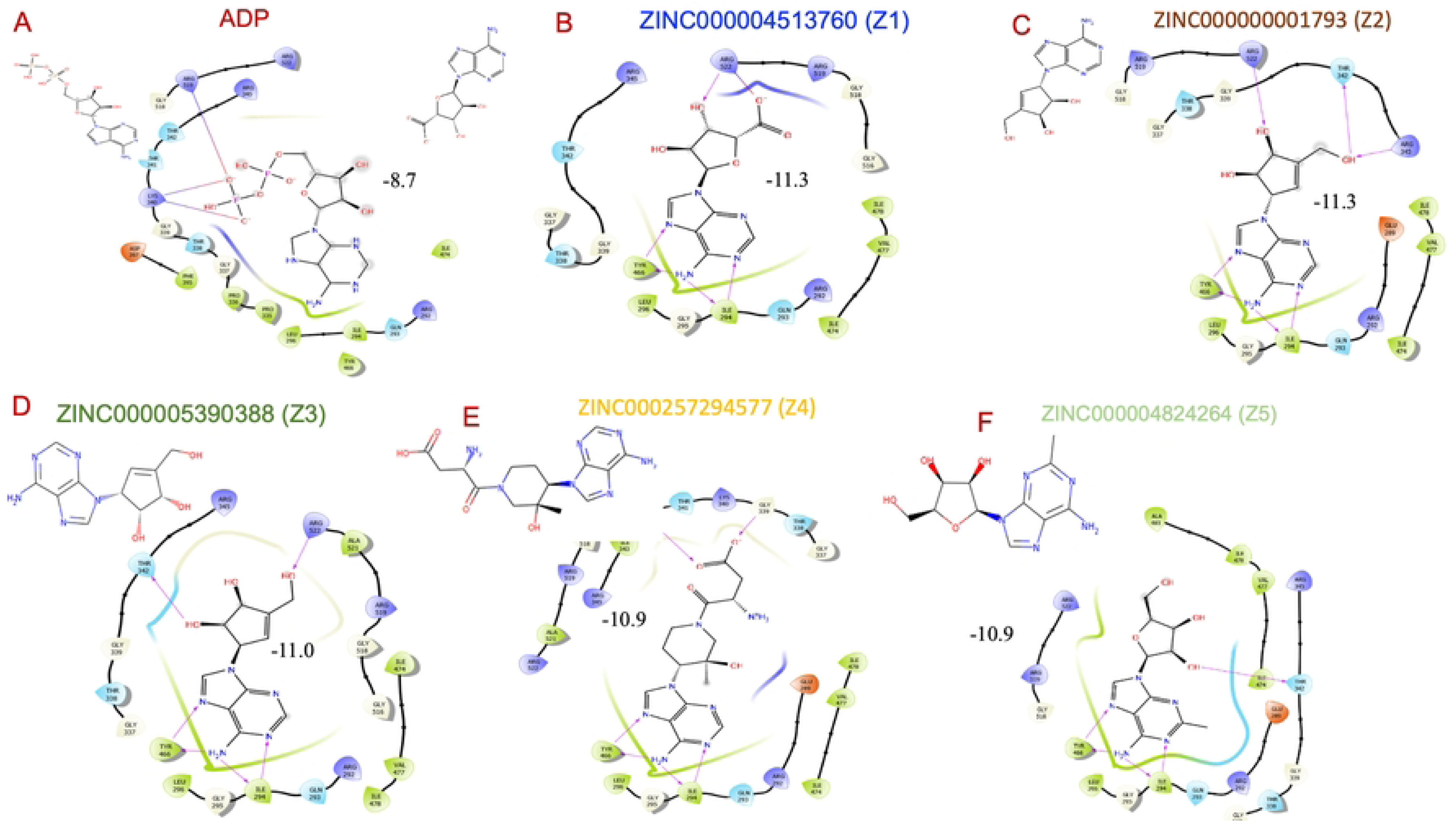
Three-dimensional view of the binding poses of the ADP and Z1-Z5 compounds at the ATPase pocket of the EccA1. **(A)** Interaction profile of the ADP with EccA1 showing key hydrogen bonding and hydrophobic interactions. **(B)** Interaction profile of the Z1 inhibitor with EccA1 showing key hydrogen bonding and hydrophobic interactions. **(C)** Interaction profile of the Z2 inhibitor with EccA1 showing key hydrogen bonding and hydrophobic interactions. **(D)** Interaction profile of the Z3 inhibitor with EccA1 showing key hydrogen bonding and hydrophobic interactions. **(E)** Interaction profile of the Z4 inhibitor with EccA1 showing key hydrogen bonding and hydrophobic interactions. **(F)** Interaction profile of the Z5 inhibitor with EccA1 showing key hydrogen bonding and hydrophobic interactions

**Table 1.**
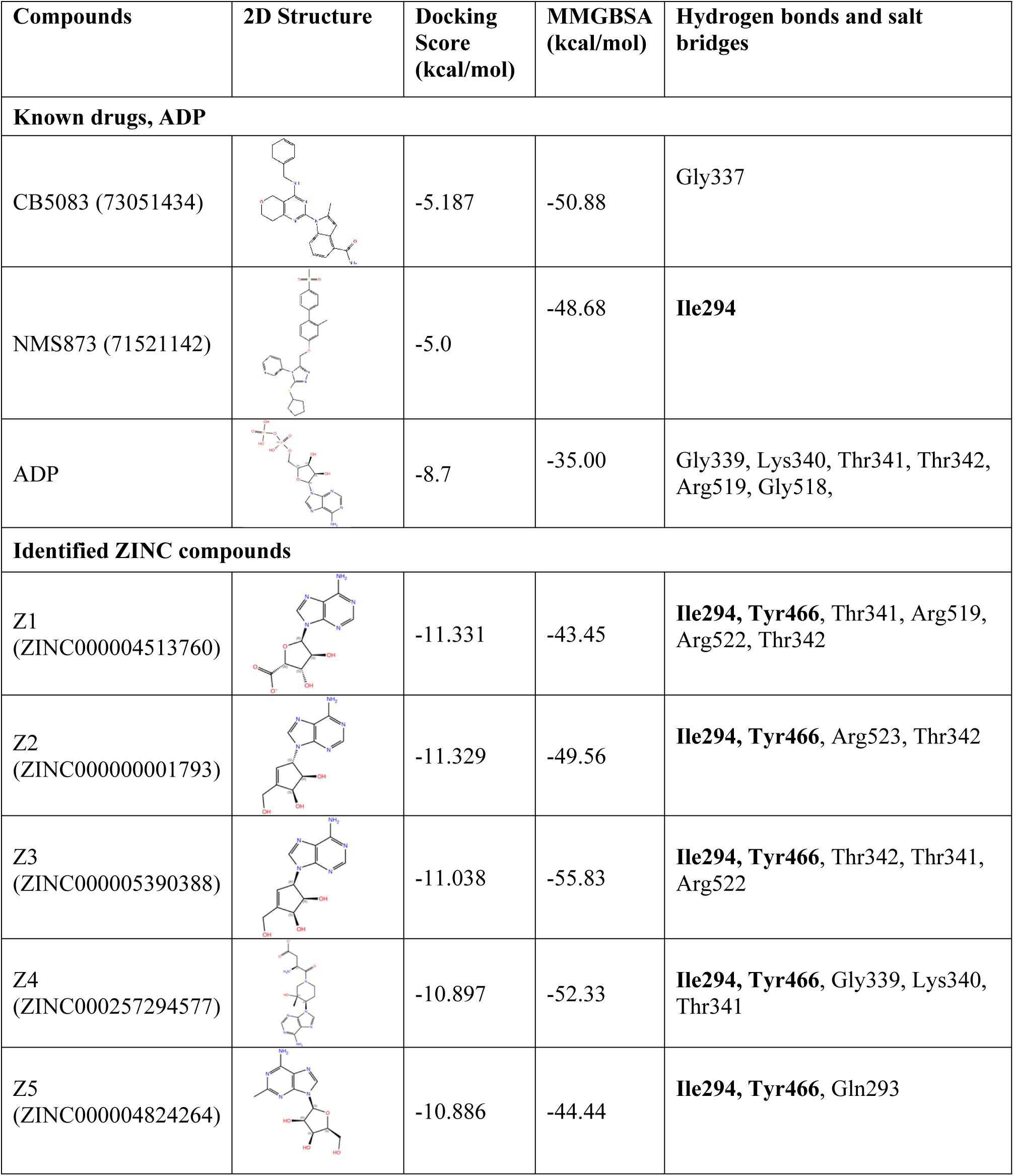
Molecular docking and binding analysis of the CB5083 and NMS873 drugs, ADP substrate, Z1-Z5 inhibitors. The two-dimensional structure, docking score, MMGBSA score and hydrogen bonds and salt bridges involve in the EccA1 recognition are shown in the table.

| Compounds | 2D Structure | Docking Score (kcal/mol) | MMGBSA (kcal/mol) | Hydrogen bonds and salt bridges |
| --- | --- | --- | --- | --- |
| **Known drugs, ADP** |  |  |  |  |
| CB5083 (73051434) |  | -5.187 | -50.88 | Gly337 |
| NMS873 (71521142) |  | -5.0 | -48.68 | Ile294 |
| ADP |  | -8.7 | -35.00 | Gly339, Lys340, Thr341, Thr342, Arg519, Gly518, |
| **Identified ZINC compounds** |  |  |  |  |
| Z1<br>(ZINC000004513760) |  | -11.331 | -43.45 | Ile294, Tyr466, Thr341, Arg519, Arg522, Thr342 |
| Z2<br>(ZINC000000001793) |  | -11.329 | -49.56 | Ile294, Tyr466, Arg523, Thr342 |
| Z3<br>(ZINC000005390388) |  | -11.038 | -55.83 | Ile294, Tyr466, Thr342, Thr341, Arg522 |
| Z4<br>(ZINC000257294577) |  | -10.897 | -52.33 | Ile294, Tyr466, Gly339, Lys340, Thr341 |
| Z5<br>(ZINC000004824264) |  | -10.886 | -44.44 | Ile294, Tyr466, Gln293 |

Maestro analysis on EccA1-ADP complex showed that Gly339, Lys340, Thr341, Thr342 residues of the Walker A motif and Arg519 residue form hydrogen bonds with PO_4_ moiety of ADP (Fig. 2A). In the EccA1-Z1 complex, purine nucleoside moiety of Z1 forms hydrogen bonds with Tyr466 and Ile294 residues and Thr341, Arg519, Arg522 and Thr342 residues of the of the EccA1 (Fig. 2B). In EccA1-Z2 complex (Fig. 2C), Ile294 and Tyr466 form hydrogen bonds with purine nucleoside moiety and Arg523 and Thr342 form hydrogen bond with the Z2 compound. In the EccA1-Z3 complex (Fig. 2D), Ile294 and Tyr466 form hydrogen bonds with the purine nucleoside moiety, and Thr341, Thr342, and Arg522 form trifurcated hydrogen bonds with the Z3 compound. In the EccA1-Z4 complex (Fig. 2E), Ile294 and Tyr466 form hydrogen bonds with the purine nucleoside moiety, and Gly339, Lys340, and Thr341 form trifurcated hydrogen bonds with the Z4 compound. In the EccA1-Z5 complex (Fig. 3F), Ile294 and Tyr466 form hydrogen bonds with the purine nucleoside moiety, and Gln293 forms hydrogen bonds with Z5 compound.

**Fig 3.**
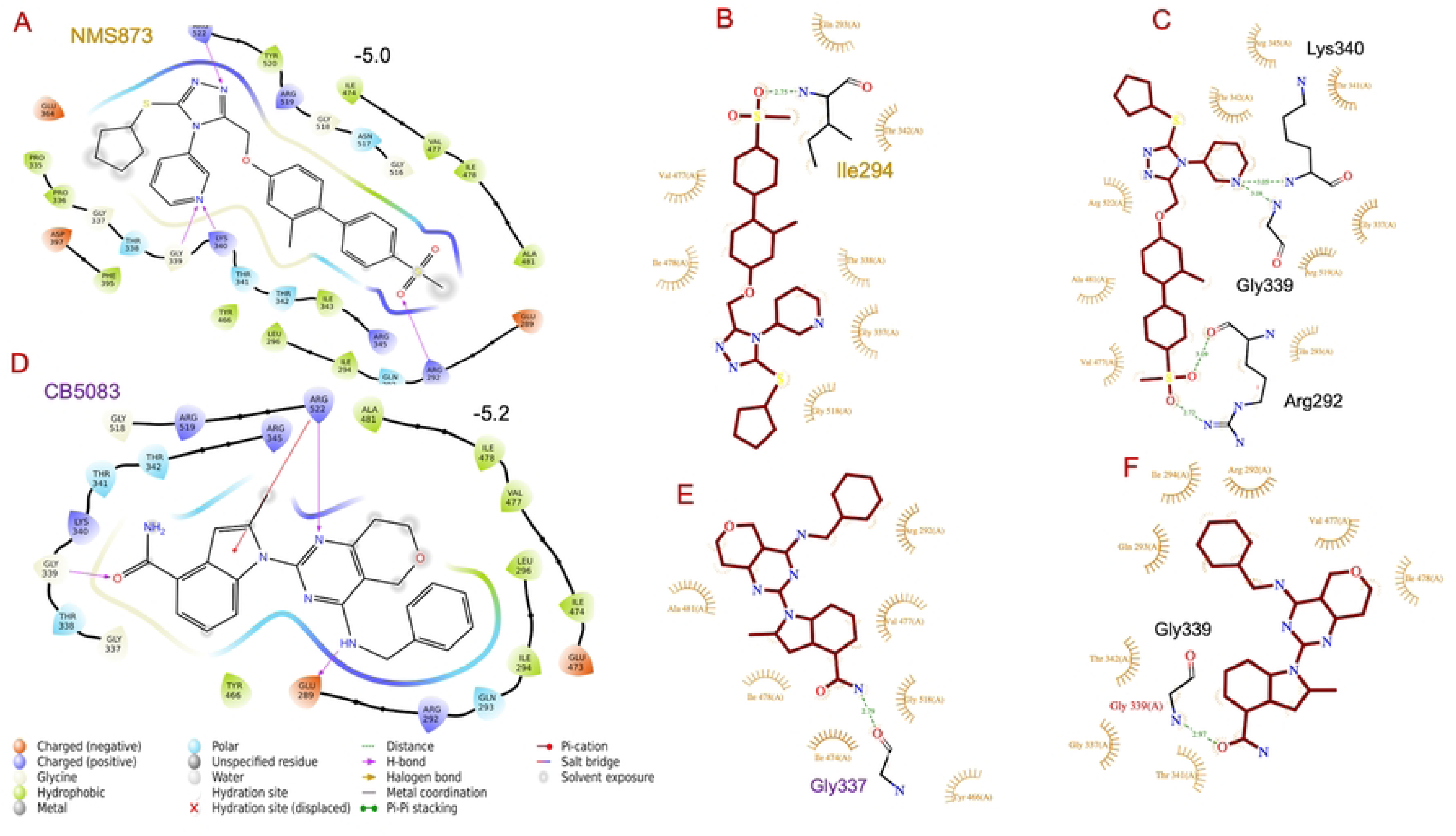
Three-dimensional view of the binding poses of the NMS-873 and CB-5083 drugs at the ATPase pocket of the EccA1. **(A)** Interaction profile of the NMS873 inhibitor with EccA1 showing key hydrogen bonding and hydrophobic interactions. **(B)** LigPlot analysis of the EccA1-NMS873 complex obtained after 100 ns dynamics simulation. **(C)** LigPlot analysis of the EccA1-NMS873 complex obtained at 0 ns dynamics simulation. **(D)** Interaction profile of the CB5083 inhibitor with EccA1 showing key hydrogen bonding and hydrophobic interactions. **(E)** LigPlot analysis of the EccA1-CB5083 complex obtained after 100 ns dynamics simulation. **(F)** LigPlot analysis of the EccA1-CB5083 complex at 0 ns dynamics simulation.

### Molecular docking and interactions analysis of the CB5083 and NMS873 drugs

Glide docking module of Schrödinger [10] was employed with advanced algorithms to predict the binding affinity and geometry of the CB5083 and NMS873 drugs (Fig. 3A-F). A docking score of -5.2 kcal/mol and binding affinity -50.88 kcal/mol was observed for the CB5083 drug against EccA1. LigPlot [35] analysis showed that Gly339(N) of Walker A motif forms a hydrogen bond with the CB5083 drug. In addition, other residues of the ATPase signature motif Gly334-Thr342 and Gly292-Leu296 region of the EccA1 were involved in the van der Waals interactions with CB5083 drug. In the earlier p97-CB5083 complex [36], the drug competes for ATP binding at the ATPase pocket of p97 ATPase.

In the case of the EccA1-CB5083 drug complex, only Gly339 formed a hydrogen bond with the CB5083 drug. After a 100 ns simulation, Gly337 was forming a hydrogen bond with CB5083. In the case of EccA1-NMS873 drug complex, Lys340 and Gly339 formed hydrogen bonds with one end, and Arg292 formed at the other end. After a 100 ns simulation, only Ile294 formed a hydrogen bond to the NMS873 drug. (Fig. 3A-F).

The EccA1 docking with the NMS873 drug yielded a docking score of −5.0 kcal/mol and a binding affinity of 48.68 kcal/mol. LigPlot [35] analysis showed that Gly339(N) and Lys340(N) of the Walker A motif and Arg292(N, Nε) form a hydrogen bond with the NMS873 drug. In addition, other residues of the ATPase signature motif Gly334-Thr342 form van der Waals interactions with the NMS873 (Fig. 3A-F). In the earlier p97-NMS873 complex [37], NMS873 covalently modifies the Cys523 of the ATPase signature motif of the p97 and blocks the ATP binding.

### Molecular dynamics simulation of the Apo EccA1 and its complexes with (i) ADP, (ii) Z1-Z5 inhibitors, (iii) CB5083 and NMS873 drugs

A 100 ns dynamics simulation was performed on apo EccA1 and its complexes with ADP, Z1-Z5 compounds, CB5083, and NMS873 drugs using the Desmond module of Schrödinger [28]. The trajectories were analyzed and calculated the RMSD, Rg, RMSF, hydrogen bonds, solvent accessible surface area, and molecular interactions using the simulation interaction diagram wizard (SID) of the Desmond program.

LigPlot analysis on the EccA1 complexed with ADP, Z1-Z5 compounds, CB5083, and NMS873 drugs was performed at 0 and 100 ns simulation and analyzed the conformational changes due to ligand binding. In the EccA1-ADP complex, Gly339, Lys340, Thr341, Arg519 residues of EccA1 formed hydrogen bonds with the di-PO_4_ moiety of the ADP. Noo hydrogen bond was formed with the adenine ring of the ADP (Fig. 4A, black letters). After a 100 ns simulation, the Lys340, Thr341, Thr342, and Arg519, still formed hydrogen bonds with the di-PO_4_ moiety of the ADP; however, new Arg522, Ile294, and Tyr466 residues appeared to form hydrogen bonds with the adenine ring (Fig. 4A, red letters). These data showed that the Adenine ring fitted better in the ATPase groove after 100 ns simulation and Adenine ring gained hydrogen bonding from other residues of EccA1.

**Fig. 4.**
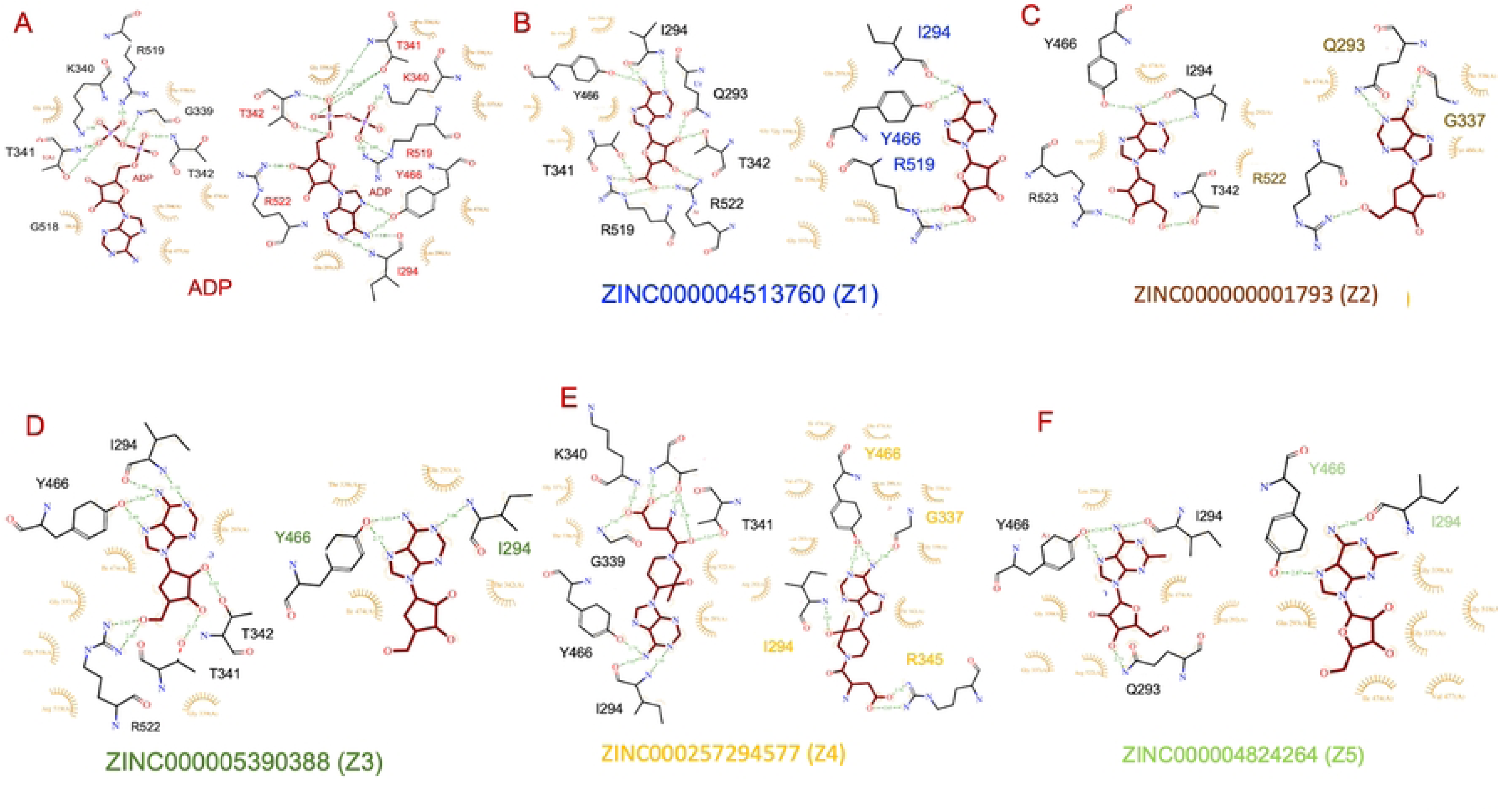
LigPlot analysis of the EccA1 complexed with ADP and Z1-Z5 inhibitors after 100 ns dynamics simulation. **(A)** LigPlot analysis of simulated EccA1+ADP complex (red letters) and starting EccA1+ADP complex (black letter). **(B)** LigPlot analysis of simulated EccA1+Z1 complex (maroon letters) and starting EccA1+ADP complex (black letter). **(C)** LigPlot analysis of simulated EccA1+Z2 complex (yellow letters) and starting EccA1+Z2 complex (black letter). **(D)** LigPlot analysis of simulated EccA1+Z3 complex (cyan letters) and starting EccA1+Z3 complex (black letter). **(E)** LigPlot analysis of simulated EccA1+Z4 complex (blue letters) and starting EccA1+Z4 complex (black letter). **(F)** LigPlot analysis of simulated EccA1+Z5 complex (letters) and starting EccA1+Z5 complex (black letter).

In the EccA1-Z1 complex, the Adenine ring formed hydrogen bonds with Tyr466, Ile294, Gln293, and Thr342 of EccA1. The Arg522, Arg519, and Thr341 residues of the EccA1 formed additional hydrogen bonds with the phosphate ring of the Z1 inhibitor (Fig. 4B, black letter). After a 100 ns simulation, Tyr466 and Ile294 residues formed hydrogen bonds with Adenine ring, and only Arg519 formed a hydrogen bond with the ribose sugar ring of the Z1 inhibitor (Fig. 4B, blue letter). In the EccA1-Z2 inhibitor complex, Tyr466 and Ile294 residues formed hydrogen bonds with Adenine, and Arg519 formed a hydrogen bond with the ribose sugar ring of the Z2 inhibitor (Fig. 4C, black letter). After 100 ns simulation, Gly337 and Gln293 formed hydrogen bonds with the Adenine ring, and Arg522 formed a hydrogen bond with the ribose sugar ring of the Z2 inhibitor (Fig. 4C, orange letter). In the EccA1-Z3 inhibitor complex, Tyr466 and Ile294 residues formed hydrogen bonds with Adenine and Thr341, Thr342 and Arg522 formed hydrogen bonds with ribose sugar ring of the Z3 inhibitor (Fig. 4D, black letter). After a 100 ns simulation, Tyr466 and Ile294 residues were involved in hydrogen bonding with Adenine. No hydrogen bond was observed with the ribose sugar ring of Z3 inhibitor (Fig. 4D, green letter).

In the EccA1-Z4 inhibitor complex, Tyr466 and Ile294 residues formed hydrogen bonds with Adenine and Gly339, Lys340 and Thr341 formed hydrogen bonds with the ribose sugar ring of the Z4 inhibitor (Fig. 4E, black letter).. After 100 ns simulation, Tyr466 and Gly337 formed hydrogen bonds with the Adenine ring and Ile294, and Arg345 formed hydrogen bonds with the ribose sugar ring of the Z4 inhibitor (Fig. 4D, yellow letter). In the EccA1-Z5 inhibitor complex, Tyr466 and Ile294 residues formed hydrogen bonds with the Adenine ring, and Gln293 formed a hydrogen bond with the ribose sugar ring of Z5 inhibitor (Fig. 4F, black letter).After a 100 ns simulation, Tyr466 and Ile294 residues formed hydrogen bonds with the Adenine ring, and no hydrogen bond was observed with the ribose sugar ring of Z5 inhibitor (Fig. 4F, light green letter).

Together, these data indicated that Z1-Z5 inhibitors fitted well into the ATPase pocket of EccA1, involved in multiple hydrogen bonding with ATPase site residues and formed stable complex. The CB5083 and NMS873 drugs against p97 ATPase did not fitted well into EccA1 ATPase pocket like ADP substrate and Z1-Z5 compounds.

### Dynamics simulation parameters analysis of Apo EccA1 and its complexes with (i) ADP (ii) Z1-Z5 inhibitors and (iii) CB-5083 and NMS-873 drugs

#### Root mean square deviation (RMSD) analysis

The RMSD plot on apo and complexed EccA1 showed stable structures and minor conformation changes during 100 ns simulations (Fig. 5A). For apo EccA1, the RMSD varies from ∼2 - 20 Å during the first 60 ns simulation and remains stable to ∼17 Å during the remaining 40 ns simulation. For EccA1-ADP complex, RMSD started from ∼ 2 Å and reached ∼ 15 Å at 40 ns simulation, and remained stable at ∼12 Å during another 60 ns simulation. It indicated that ADP binding was a little unstable till 20 ns simulation and stabilized during the remaining 80 ns simulation. It showed that ADP binding to EccA1 has enhanced the stability of the EccA1 complex. In the EccA1-Z1 complex, RMSD increased ∼2-12 Å during the first 10 ns simulation and further increased to 20 Å during the 60 ns simulation. The RMSD decreased to ∼15 Å during 60-100 ns simulation to attain a low-energy structure. In the EccA1-Z2 complex, the RMSD increased ∼3-9 Å during the first 10 ns simulation and fluctuates to ∼17–19 Å during the remaining 90 ns simulation. In the EccA1-Z3 complex, RMSD increased to ∼ 2-15 Å during the first 10 ns simulation and remained stable at ∼ 15-17 Å during the 10-90 ns simulation. In the EccA1-Z4 complex, initially 10 ns simulation, the RMSD increased to 2-12 Å and fluctuated between 8-12 Å during 10-50 ns simulation. During 50-100 ns, the RMSD decreased to ∼11-7 Å. In the EccA1-Z5 complex, the RMSD increased ∼2-13 Å in the first 10 ns simulation and increased to ∼6-10 Å during 10-100 ns simulation. In EccA1-NMS873 drug complex, the RMSD increased to ∼2-10 Å during 0-15 ns simulation and stabilized to ∼7-8 Å during 15-100 ns simulation. In case of EccA1-CB5083 complex, the RMSD increased to ∼3-11 Å in 0-15 ns simulation and remained stable to ∼9-10 Å during 15-100 ns simulation.

**Fig. 5.**
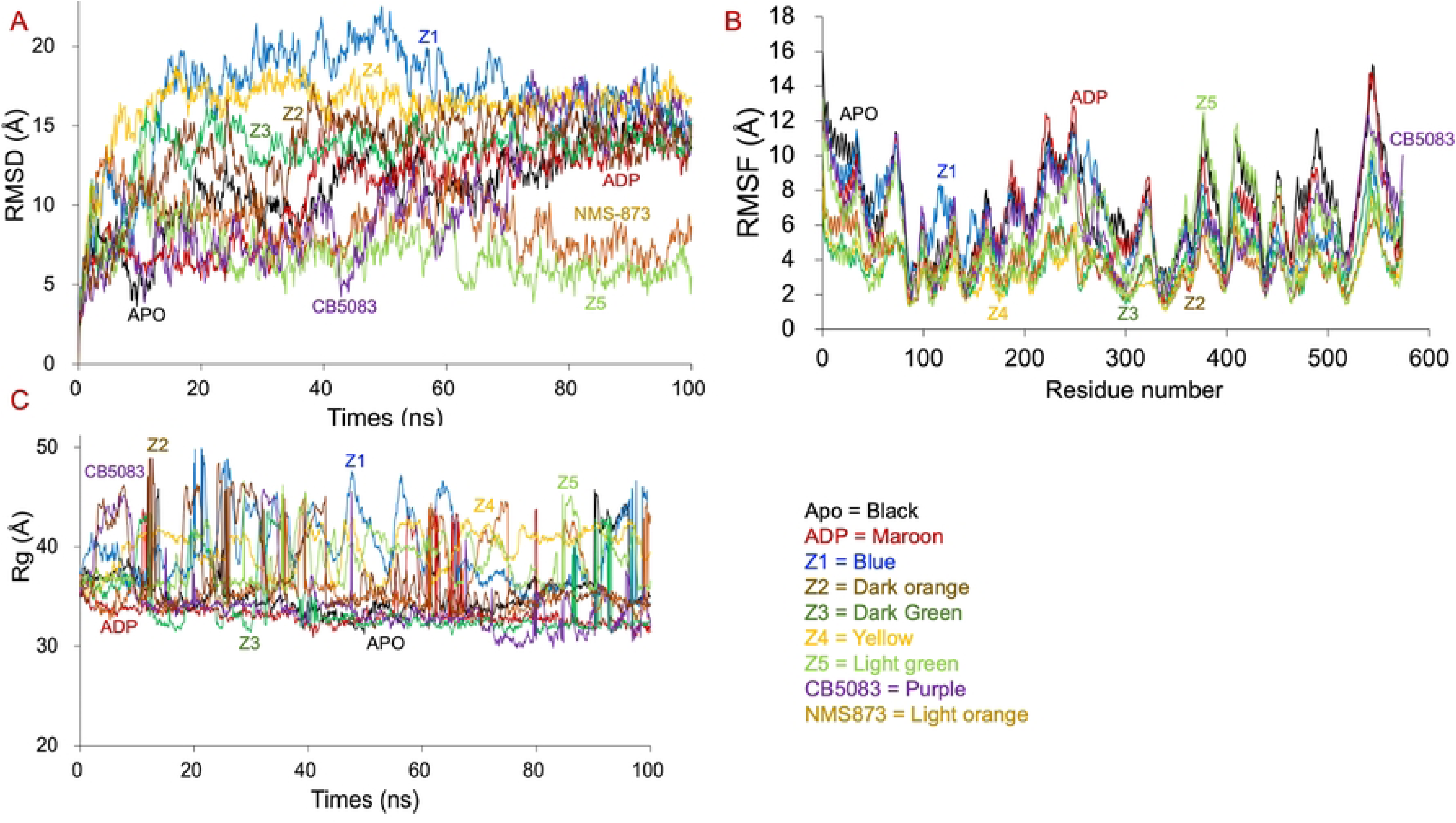
Structural dynamics simulation of the apo EccA1 (black) and its complexes with (i) ADP (iii) Z1 (iv) Z2 (v) Z3 (vi) Z4 (vii) Z5 (viii) CB5083 and (vii) NMS873 during 100 ns simulation. **(A)** The Root-mean-square deviation (RMSD) plot of the apo EccA1 (black) and its complexes with (i) ADP (maroon) (iii) Z1(blue) (iv) Z2 (dark orange) (v) Z3 (dark green) (vi) Z4 (yellow) (vii) Z5 (light green) (viii) CB5083 (purple) and (vii) NMS873 (light orange) after 100 ns simulation. **(B**) The Root-mean-square fluctuation (RMSF) plot of the apo EccA1 (black) and its complexes with (i) ADP (maroon) (iii) Z1(blue) (iv) Z2 (dark orange) (v) Z3 (dark green) (vi) Z4 (yellow) (vii) Z5 (light green) (viii) CB5083 (purple) and (vii) NMS873 (light orange) after 100 ns simulation. **(C)** The Radius of Gyration (Rg) plot of the apo EccA1 (black) and its complexes with (i) ADP (maroon) (iii) Z1(blue) (iv) Z2 (dark orange) (v) Z3 (dark green) (vi) Z4 (yellow) (vii) Z5 (light green) (viii) CB5083 (purple) and (vii) NMS873 (light orange) after 100 ns simulation.

The Z1 to Z5 ligands remained stable in the ATPase pocket and did not dissociate during 100 ns simulation. The ADP and Z5 ligands showed the most stable binding compared to the Z1 to Z4 ligands. ADP and Z5 both reduce flexibility of the binding-site residues and formed effective binding with EccA1. The apo EccA1 showed higher flexibility during the 100 ns simulation compared to complexed EccA1. The Z5 compound showed most stable complex with EccA1, as indicated by most stable RMSD profile. The Z4 compound showed moderate stability compared to Z1-Z3 compounds. The CB5083 and NMS873 exhibited the largest RMSD fluctuation during 100 ns simulation compared to the ADP and Z1-Z5 compounds. The ADP is a natural substrate that showed most stable binding during RMSD analysis.

#### Radius of gyration analysis

Rg was calculated for apo EccA1 and its complexes with (i) ADP, (ii) Z1-Z5 compounds, and (iii) CB5083, NMS-873 drugs during a 100 ns simulation (Fig. 5B). The Rg measures the overall compactness of the structure. ∼35-45 Å fluctuation in Rg for apo EccA1 and ∼32-36 Å fluctuation for EccA1-ADP complex were found during 100 ns simulation. These data indicated that EccA1 stability enhance because of ADP binding. In EccA1-Z1 complex, ∼35-47 Å fluctuation during 100 ns simulation, ∼35-43 Å for the EccA1-Z2 complex, ∼32-44 Å for EccA1-Z3 complex, ∼35-43 Å in case of EccA1-Z2 complex, ∼32–44 Å in case of EccA1-Z3 complex, ∼35–45 Å for EccA1-Z4 complex and ∼32–44 Å for EccA1-Z3 complex and ∼32–43 Å for EccA1-Z5 complex were observed. In case of CB5083-EccA1 complex, ∼45 Å fluctuation during 0-20 ns simulation and 33-35 Å during 20-100 ns simulation were observed. In case of the EccA1-NMS873 complex, ∼35-48 Å fluctuation during 0-45 ns simulation and ∼33-36 Å fluctuation during 45-100 ns simulation were observed.

The Rg profile indicated that ADP formed the most stable complex with EccA1. The Z1-Z5 inhibitors also formed a stable complex like ADP. The CB5083 and NMS873 drugs showed significant fluctuations in the beginning and stabilized during 100 ns simulation. However, they did not form a stable complexed structure with EccA1.

#### Root mean square fluctuation analysis

RMSF was calculated for apo EccA1 and its complexes with ADP, Z1-Z5 compounds, CB5083, and NMS-873 drugs during 100 ns simulation (Fig. 5C). In Apo EccA1 (black), Asp540-Asp547 residues showed maximum RMSF (6-14 Å), Gly376 (12.1 Å), and Met1-Ser7 (4-12 Å) during 100 ns simulation. In the EccA1-ADP complex (marron), the Val539-Glu547 showed reduced RMSF (4-12 Å), and Gly220-Ser249 showed RMSF (4-12 Å) during 100 ns simulation. These data showed that ADP stabilized the overall EccA1 structure. Among all complexes, the EccA1 complexed with Z1(blue) and Z5 (light green) inhibitors showed higher RMSF (3-12 Å), while Z4 (yellow), Z3 (dark green), and Z2 (dark orange) inhibitors showed lower RMSF (2-10 Å). The EccA1-CB5083 complex (purple) showed the RMSF (4-10 Å), and EccA1-NMS873 (light orange) showed the RMSF (4-10 Å) during a 100 ns simulation. Maximum RMSF was observed for Asp540-Asp547, Gly220-Ser249, and Met1-Ser7 residues. The EccA1-Z4 inhibitor showed the lowest RMSF across all complexes and EccA1-ADP complex showed lower RMSF fluctuations compared to the Apo EccA1 structure. The Z1, Z5, CB5083, and NMS873 complexes showed higher fluctuations, indicating weaker binding.

#### SASA analysis

SASA analysis was performed on EccA1 complexed with (i) ADP, (ii) Z1-Z5 compounds, (iii) CB5083 and NMS873 drugs during a 100 ns simulation. Lower SASA values showed a more compact and stable structure (more compact surface), and higher SASA values showed higher flexibility and unstable structure (more exposed surface) (Fig. 6A).

**Fig. 6.**
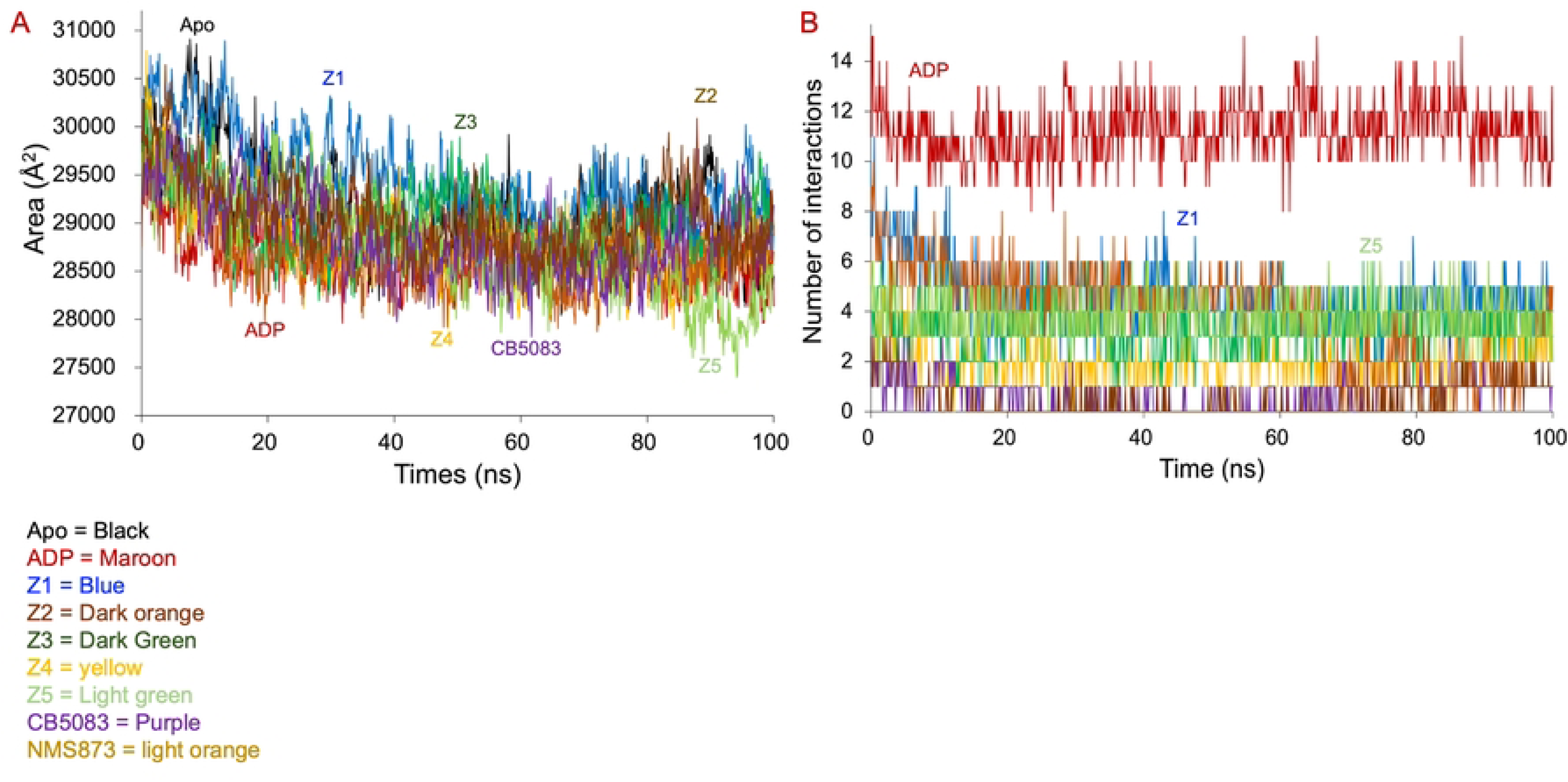
Intermolecular hydrogen bonds and SASA analysis of the apo EccA1 (black) and its complexes with (i) ADP (iii) Z1 (iv) Z2 (v) Z3 (vi) Z4 (vii) Z5 (viii) CB5083 and (vii) NMS873 during 100 ns dynamics simulation. **(A)** Hydrogen bond analysis of apo EccA1 (black) and its complexes with (i) ADP (maroon) (iii) Z1(blue) (iv) Z2 (dark orange) (v) Z3 (dark green) (vi) Z4 (yellow) (vii) Z5 (light green) (viii) CB5083 (purple) and (vii) NMS873 (light orange) after 100 ns simulation. **(B)** The solvent accessible surface area (SASA) of the apo EccA1 (black) and its complexes with (i) ADP (maroon) (iii) Z1(blue) (iv) Z2 (dark orange) (v) Z3 (dark green) (vi) Z4 (yellow) (vii) Z5 (light green) (viii) CB5083 (purple) and (vii) NMS873 (light orange) after 100 ns simulation.

For apo EccA1, the SASA starts from ∼30,500 Å² and decreases to ∼28,500 Å², indicating large fluctuations, structural flexibility, and rearrangements without ligand. In the EccA1-ADP complex, the SASA peak stabilizes around ∼29,000 Å² quickly, indicating that ADP binding induces a more compact and stable conformation to EccA1. In the EccA1-Z1 inhibitor complex, the SASA steadily decreased from ∼30,500 Å² to ∼29,000 Å², indicating structural compactness over time. In the EccA1-Z2 inhibitor complex, the SASA peak dropped early and stabilized around ∼28,700–29,000 Å², like ADP. In the EccA1-Z3 complex, the SASA started around ∼29,500 Å² and fluctuates throughout the 100 ns simulation. It exhibited moderate stability, but higher surface exposure compared to the EccA1-ADP and EccA1-Z1 complex. In the EccA1-Z4 complex, the SASA rapidly decreased and stabilized at ∼28,800 Å². These data indicated a strong and compact interaction with EccA1. In the EccA1-Z5 complex, the SASA declined steadily to ∼28,000 Å², indicating the most compact conformation of the complex. The Z5 compound showed the most significant reduction in SASA and the most stable compact structure. The ADP and Z1-Z4 compounds also contribute significantly to protein compaction. The apo EccA1 showed the highest fluctuation, indicating a flexible structure without ligand.

In the EccA1-CB5083 complex, SASA fluctuates 29,800 to 30,000 Å² during 0-40 ns simulation and remained stable during 40 to 100 ns simulation. In the EccA1-NMS873 complex, the SASA starts from ∼30,000 Å² and gradually decreased to ∼29,000 Å² during the 0-40 ns simulation. During the 40-100 ns simulation, it remained 30,000 Å², indicating transient expansions in surface exposure. NMS873 exhibited a SASA pattern like apo EccA1; however, CB5083 fluctuated before reaching a slightly lower SASA value.

The SASA values of all complexes decreased compared to apo EccA1. ADP showed a similar value to that of the apo EccA1 enzyme. The Z5 complex showed a stable and consistent SASA profile better than other ligands. Z1, Z2, Z3, and Z4 compounds showed moderate stability during the 100 ns simulation; CB5083 and MNS873 exhibited higher fluctuations (weak structural stability).

#### Hydrogen bonds analysis

During a 100 ns simulation, hydrogen bond analysis was performed on EccA1 complexed with ADP, Z1-Z5 inhibitors, and CB5083 and NMS873 drugs (Fig. 6B). In the EccA1-ADP complex, ∼ 8-16 hydrogen bonds were formed during 100 ns simulation, indicating high stability. In the EccA1-Z1 inhibitor complex, ∼ 2-10 hydrogen bonds were formed during a 100 ns simulation. Initially, ∼ 8-10 hydrogen bonds were formed, and decreased to ∼4-6 hydrogen bonds after a 20 ns simulation. In the EccA1-Z2 inhibitor complex, ∼0-5 hydrogen bonds were formed in the beginning and later decreased to 0-2 hydrogen bonds, which indicated the poor binding and stability. In the EccA1-Z3 inhibitor complex, ∼0-6 hydrogen bonds were formed in the beginning, better than the EccA1-Z2 complex. In the EccA1-Z4 inhibitor complex, ∼8-10 hydrogen bonds formed in the beginning and decreased to ∼2-4 after 60 ns simulation. In the EccA1-Z5 inhibitor complex, ∼1-6 hydrogen bonds were formed, and ∼3-5 hydrogen bonds were observed during a 100 ns simulation.

In the EccA1-CB5083 drug complex, ∼1-3 hydrogen bonds formed during 0-15 ns, and ∼2 hydrogen bonds were formed during 15-100 ns of dynamics simulation. It indicated that the EccA1-CB5083 complex showed higher fluctuations during 0-15 ns simulation and attained stability during 15-100 ns simulation. In EccA1-NMS873 complex, ∼0-3 hydrogen bonds were observed during 0-10 ns simulation and maintained 1 hydrogen bond during 10-100 ns simulation. During 0-10 ns, the NMS873 occupies the binding pocket and did not form a stable complex. During 10-100 ns, the NMS873 drug attained moderate stability in the binding pocket. The CB5083 and NMS873 drugs bind strongly to EccA1 and remain stable.

The ADP showed the highest hydrogen-bond count during the 100 ns simulation and binds strongly to EccA1. The CB5083 and NMS873 exhibited a weak binding profile and were characterized by few Hydrogen bonds. In Z1-Z5 compounds, Z5 showed the best and most stable interaction, a consistent hydrogen bond profile, and less fluctuations compared to Z1-Z4 compounds. The Z1 compound exhibited moderate fluctuations, starting from 8–9 hydrogen bonds and remained stable to 4–5 hydrogen bonds, better than Z2 to Z4 compounds.

#### ADMET properties and Lipinski’s rule analysis on the Z1-Z5 inhibitors

Absorption, Distribution, Metabolism, and Excretion (Swiss-ADME) properties of identified Z1-Z5 compounds were calculated to assess the bioavailability, therapeutic effectiveness, safety, and dosages of these inhibitors. These data provide information about how the body handles the drugs, contribute to drug designing, regulatory approval, and personalized treatment strategies. Lipinski’s rule of five was used for the identification of the goodness for oral drug administration in Z1-Z5 compounds. Following criteria: (i) Mw∼500 Da or less (ii) lipophilicity (LogP) ∼ 5 or less and (iii) number of the hydrogen bond acceptors (≤ 15) and donors (≤ 5). The QPlogPo/w value between 5 and 45 indicated high solubility in the lipophilic solvents and low solubility in water.

The cellular permeability (nm/second) of ZINC compounds was determined using QPPMDCK (model for gut-blood barriers) and QPPCaco and showed values lower than 500, indicating low permeability. These values were higher for CB5083 and NMS873 drugs, indicating better permeability of these ZINC compounds. The QPlogPo/w parameter indicates that the solubility parameters for potential inhibitors ideally should lie between 5 and 45. For Z1-Z5 compounds, this parameter was found to be less than 4, indicating high solubility in the lipophilic solvents and low solubility in water. Further, the QPlogBB parameter indicated whether the compound could cross the blood-brain barrier, and negative values indicated a low transfer rate with no active transport. We have obtained less than −1.921 QPlogBB values for identified ZINC compounds. For CB5083 and NMS873 drugs, the −0.907 lowest value was obtained. The human oral consumption values for Z1-Z5 compounds were found in the range of 30-55%. All Z1-Z5 inhibitors showed predicted pharmacokinetic characteristics and obeyed Lipinski’s rule of five (Table 2) and drug-like characteristics (Table 3).

**Table 2.**
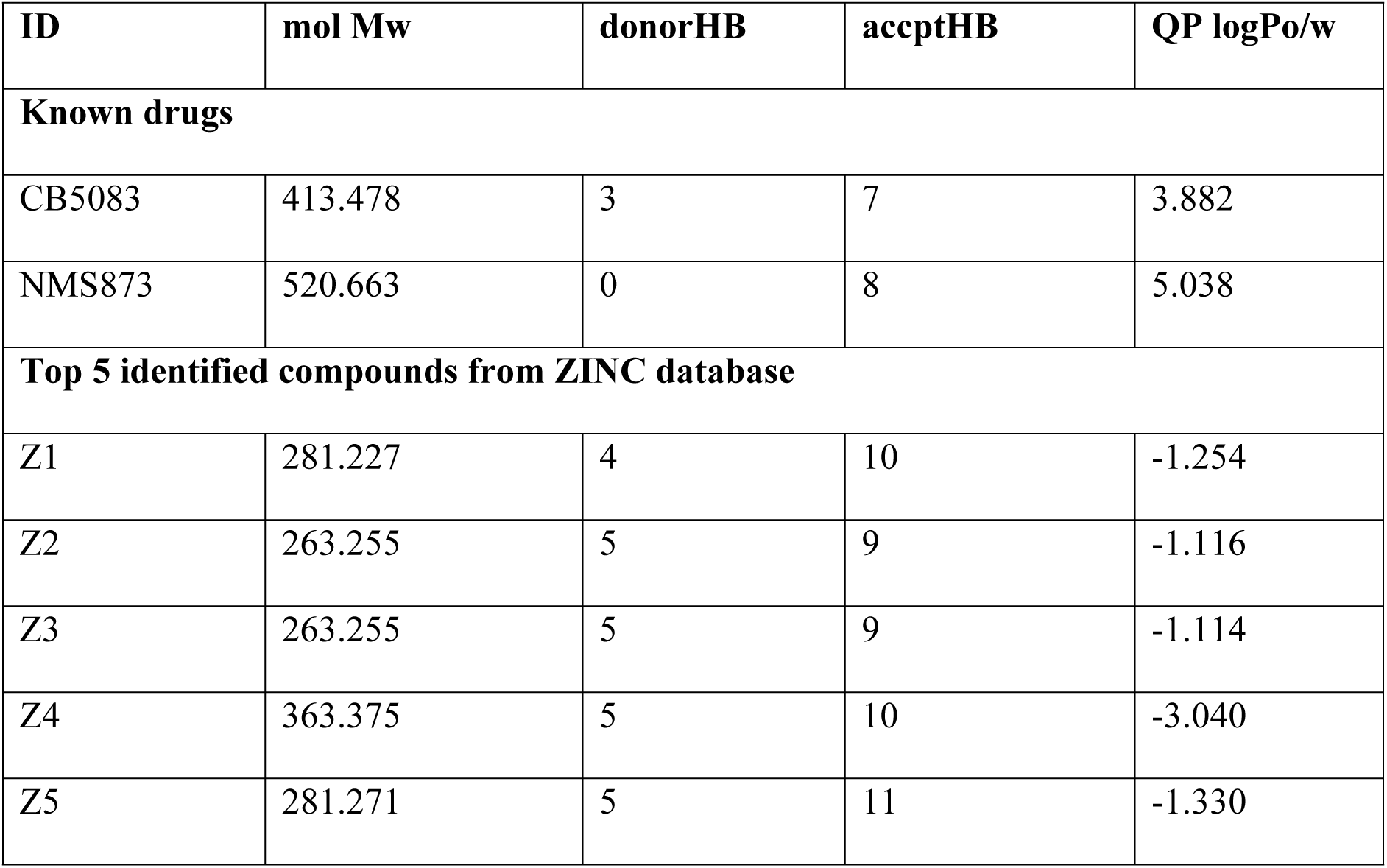
Lipinski’s Rule of Five predictions and lysis on Z1-Z5 inhibitors and two drugs.

| ID | mol Mw | donorHB | acceptHB | QP logPo/w |
| --- | --- | --- | --- | --- |
| **Known drugs** |  |  |  |  |
| CB5083 | 413.478 | 3 | 7 | 3.882 |
| NMS873 | 520.663 | 0 | 8 | 5.038 |
| **Top 5 identified compounds from ZINC database** |  |  |  |  |
| Z1 | 281.227 | 4 | 10 | -1.254 |
| Z2 | 263.255 | 5 | 9 | -1.116 |
| Z3 | 263.255 | 5 | 9 | -1.114 |
| Z4 | 363.375 | 5 | 10 | -3.040 |
| Z5 | 281.271 | 5 | 11 | -1.330 |

**Table 3.** The ADMET properties of lead ZINC compounds and two drugs, calculated using QikProp program.

| ZINC ID | Absorption | Distribution |  |  | CNS permeability |  |
| --- | --- | --- | --- | --- | --- | --- |
|  | %Human oral consumption | QPPCaco | QPPMDCK | QPlogPo/w | CNS | QPlogBB |
| **Known drugs/inhibitor** |  |  |  |  |  |  |
| CB5083 | 100.000 | 758.061 | 366.711 | 0.553 | -1 | -0.907 |
| NMS873 | 75.313 | 317.929 | 196.882 | 0.737 | -2 | -1.585 |
| **5 identified Z1-Z5 compounds** |  |  |  |  |  |  |
| Z1 | 30.807 | 4.226 | 1.709 | -0.960 | -2 | -2.003 |
| Z2 | 49.549 | 42.464 | 16.271 | -0.784 | -2 | -1.921 |
| Z3 | 50.380 | 47.184 | 18.234 | -0.782 | -2 | -1.851 |
| Z4 | 0.000 | 0.419 | 0.267 | -0.977 | -2 | -2.162 |
| Z5 | 55.106 | 101.976 | 41.942 | -0.797 | -2 | -1.406 |

## Discussion

There is an urgent need to explore and characterize new drug targets to treat multidrug-resistant, extensively drug-resistant, and totally drug-resistant tuberculosis. The ESX-1 secretion system is essential for phagosomal escape, host cell lysis, immune modulation, and full virulence of *M. tuberculosis*. The EccA1 enzyme from the ESX-1 secretion system is involved in the secretion of the ESX-1 and PE/PPE virulence factors. Disruption of the EccA1 attenuates virulence in the macrophages and animal models. ATPase pocket of the EccA1 is unique and highly conserved with no human analog. EccA1-mediated ATP hydrolysis and substrate engagement are required for efficient ESX-1 secretion, and selective inhibition will reduce intracellular survival and host cell damage by *M. tuberculosis*.

Here, an *in-silico* approach was used to identify the novel ATPase inhibitors against EccA1. All compounds from the ZINC database were screened using *in silico* docking technique, and the top five ZINC compounds were identified, having the highest binding affinity to EccA1, higher than the ADP substrate. In addition, docking and dynamics simulation analysis was performed on EccA1 complexed with ADP and NMS-873 and CBS-508 drugs, and compared the discovered ZINC compounds. The top selected Z1-Z5 compounds showed binding energy in the range (−43 to -55 kcal/mol), higher than ADP (−35.7 kcal/mol), indicating potential for promising therapeutic development. The ZINC compounds (Z1-Z5) showed high docking scores (−10.88 to -11.33 kcal/mol) compared to ADP (−8.70 kcal/mol), NMS873 (−5.00 kcal/mol), and CB5083 (−5.19 kcal/mol) drugs. These data indicated a competitive binding of discovered ZINC compounds to EccA1 ATPase pocket to inhibit the enzyme activity.

100 ns molecular dynamics simulation was performed on apo EccA1 and its complexes with ADP, Z1-Z5 compounds, CB-5083 and NMS-873 drugs. Molecular dynamics simulation on apo and ligand-bound EccA1 provides information about ligand-induced conformational changes, changes in the flexibility and stability of the protein, binding site dynamics, and allosteric effects. Molecular dynamics simulation showed significant stability of the Z1-Z5 compounds in the EccA1 ATPase pocket. The RMSD, RMSF, Rg, SASA, and hydrogen bonding profiles were calculated for Apo and ligand bound EccA1 to discover the EccA1 conformational stability and compared with CB5083 and NMS873 drugs.

The RMSD parameter indicated the overall stability of the EccA1-ligand complex during the dynamic’s simulation (Fig. 5A). The Apo EccA1 showed high RMSD fluctuation (black, ∼5-15 Å), ADP (maroon, ∼5-12 Å), Z1 (blue, ∼5-25 Å), Z2 (dark orange, ∼ 5-15 Å), Z3 (dark green, ∼5-13 Å), Z4 (yellow, ∼5-17 Å) and Z5 (light green, ∼5-6 Å). These data indicated that RMSD for Z1-Z5 inhibitors considerably stabilized without conformational changes. For NMS873 (light orange, RMSD∼5-7 Å) and CB5083 (purple, RMSD ∼5-15 Å), drugs were observed, indicating a stable conformation during 100 ns simulation.

The Rg parameter characterizes the compactness and shape of the protein. The time evolution of Rg was calculated for apo EccA1 and its complexes with ligands and obtained following data, Apo EccA1 (black, ∼35 Å), ADP (maroon, ∼35 Å), Z1 (blue, ∼ 35-48 Å), Z2 (dark orange, ∼ 35-48 Å), Z3 (dark green, ∼ 35 Å), Z4 (yellow, ∼ 35-40 Å), Z5 (light green, ∼5-6 Å), NMS-873 (light orange, ∼35 Å), CB5083 (purple, ∼35-48 Å). Except for Z1, Z2, Z4, CB5083 inhibitors, the Rg values remained constant for all other ligands during the entire 100 ns dynamics simulation.

The RMSF parameter indicated the flexibility associated with each residue during the dynamics simulation. It showed the conformational plasticity of the protein at different regions upon ligand binding. Higher RMSF corresponds to high flexibility. The Z1, Z5, CB5083, and NMS873 displayed higher fluctuations, indicating weaker binding. The Z2, Z3, and Z4 showed lower fluctuation and bind strongly to EccA1. For all inhibitors, real-time interactions with EccA1 active site residues were plotted as stacked bars (Fig. S1). ADP, Z1 ligands showed the maximum interaction fractions, while NMS873, CBS5083, Z3, Z4, and Z5 compounds showed the minimum interaction fractions.

ADMET parameters were calculated to evaluate the toxicity parameters for identified ZINC and drug compounds. QikProp and Swiss ADMET analyses indicated that all identified ZINC compounds met the criteria for hydrogen bonds acceptor/donors and did not violate Lipinski’s rule of five and had acceptable pharmacokinetic properties.

## Conclusions

In the present study, the Z1-Z5 compounds have shown significant binding affinity to the EccA1 enzyme. Molecular docking and binding free energy calculations have shown a possible binding mechanism of the Z1-Z5 compounds to the ATPase pocket of the EccA1. Key interactions involved in the binding of the Z1-Z5 compounds to the ATPase pocket of the EccA1 were identified. Molecular dynamics simulation on ApoEccA1 and its complexes with ADP substrate, Z1-Z5 inhibitors, NMS873 and CBS5083 drugs showed stability of all complexes and the molecular mechanism involved in the ligand recognition. These data were generated using in silico tools; however, require *in vivo* and *in vitro* experiments to validate their inhibitory potential. Structural optimization for improved specificity, experimental validation, and *in vitro* evaluations for therapeutic potentials are required for next-generation drug development.

## Supporting information’s

**Fig. S1.** Histogram showing the various level of interactions of the EccA1 complexes with (i) ADP (maroon) (ii) Z1 (blue) (iii) Z2 (dark orange) (iv) Z3 (dark green) (v) Z4 (yellow) (vi) Z5 (light green) (vii) CB5083 (purple) and (viii) NMS873 (light orange) during 100 ns dynamics simulation.

## Acknowledgements

The authors gratefully acknowledge the financial support from the DBT project (BT/PR44988/DRUG/134/126/2022) from India for the current research work. Partial financial support from UGC-resource networking, DST-PURSE, and UGC-SAP grants from India. Ramesh gratefully acknowledges the Senior Research Fellowship from UGC, India. The facilities/laboratories support provided by DBT BUILDER (grant no. BT/INF/22/SP45382/ 2022) and DST FIST-II [grant no. SR/FST/LSII-046/2016(C)] grants from India are gratefully acknowledged. We acknowledge the help from the CIF facility staff of SLS, JNU, India.

## Author’s contribution

**Ramesh Kumar:** Molecular docking and dynamics simulation analysis. **Ajay K. Saxena:** Writing – original draft, Validation, Supervision, Resources, Project administration, Funding acquisition, Data curation, Conceptualization.

## Data availability

Data will be available upon request.

## Abbreviation used

RMSD: Root mean square deviation
Rg: Radius of gyration
RMSF: Root mean square fluctuation
ADP: Adenosine diphosphate
MDR: Multidrug resistance
XDR: Extensive drug resistance
MDR: Multidrug resistance
ADME: Absorption, Distribution, Metabolism, and Excretion
SASA: Solvent surface area
MM-GBSA: Molecular mechanics-based generalized born surface area
MD: Molecular dynamics
TPR: Tetratricopeptide repeat
CbxX/Cfxq: 
ESAT6: 
CFP10: 
FDA: Food and drug administration
SPC: Single point charges
PBC: Periodic boundary conditions
PME: Particle-Mesh Ewald
PDB: Protein data bank
Z1: ZINC000004513760
Z2: ZINC000000001793
Z3: ZINC000005390388
Z4: ZINC000257294577
Z5: ZINC000004824264
CB5083: 1-[4-(benzylamino)-5H,7H,8H-pyrano[4,3-d]pyrimidin-2-yl]-2-methyl-1H-indole-4-carboxamide
NMS873: 3-[3-cyclopentylsulfanyl-5-[[3-methyl-4-(4-methylsulfonylphenyl)phenoxy]methyl]-1,2,4-triazol-4-yl]pyridine

